# A comparative study of genomic adaptations to low nitrogen availability in Genlisea aurea

**DOI:** 10.1101/2021.02.09.430036

**Authors:** Thibaut Goldsborough

## Abstract

*Genlisea aurea* is a carnivorous plant that grows on nitrogen-poor waterlogged sandstone plateaus and is thought to have evolved carnivory as an adaptation to very low nitrogen levels in its habitat. The carnivorous plant is also unusual for having one of the smallest genomes among flowering plants. Genomic DNA is known to have a high nitrogen content and yet, to the author’s knowledge, no published study has linked nitrogen starvation of *G. aurea* with genome size reduction. This comparative study of the carnivorous plant *G. aurea*, the model organism *Arabidopsis thaliana* (Brassicaceae) and the nitrogen fixing *Trifolium pratense* (Fabaceae) attempts to investigate whether the genome, transcriptome and proteome of *G. aurea* showed evidence of adaptations to low nitrogen availability. It was found that although *G. aurea*’s genome, CDS and non-coding DNA were much lower in nitrogen than the genome of *T. pratense* and *A. thaliana* this was solely due to the length of the genome, CDS and non-coding sequences rather than the composition of these sequences.

## Introduction

*Genlisea aurea* (Lentibulariaceae) is a carnivorous plant found in Brazil that grows on waterlogged sandstone plateaus. It is thought to have evolved carnivory as an adaptation to very low nitrogen levels in its habitat (Müller K. *et al*. 2004). *G. aurea* is also unusual for having one of the smallest genomes among flowering plants with a genome length of just 43.4 Mb, resulting from a process called genome reduction in which intergenic regions and duplicated genes are removed (Leushkin, E.V. *et al*. 2013). The tiny genome of another carnivorous plant *Utricularia gibba* (Lentibulariaceae) shows that *G. aurea* is not the only carnivorous plants that grows in nitrogen poor habitats to have undergone genome size reduction (Ibarra-Laclette, E. *et al*. 2013). Despite DNA having a high nitrogen content, to the author’s knowledge, no published study has linked nitrogen starvation of *G. aurea* with genome size reduction. This project investigates whether the genome, transcriptome and proteome of *G. aurea* show evidence of adaptations to low nitrogen availability. In this study, the genome of *G. aurea* is compared to the model organism *Arabidopsis thaliana* (Brassicaceae) and to the nitrogen fixing *Trifolium pratense* (Fabaceae). *T. pratense* is known to fix nitrogen gas (N_2_) from the atmosphere with the help of nitrogen-fixing bacteria found in its roots (Davey A.G. *et al*. 1989). For this reason, *T. pratense* was taken as a control for a plant that is not nitrogen deprived. Reduction of nitrogen usage in proteomes has already been recorded when comparing plant proteins and animal proteins. Plants are generally regarded as nitrogen limited in comparison with animals and one study found a 7.1 % reduction in nitrogen use in amino acid side chains of plant proteins compared to animal proteins (Acquisti, C., Kumar and S., Elser, J.J. 2009). Another study found that parasitic microorganisms showed altered codon usage and genome composition as a response to nitrogen limitations (Seward, E.A. and Kelly, S. (2016).

## Methods

All genomic data was obtained from the NCBI genome database Genbank (available at ftp://ftp.ncbi.nlm.nih.gov/genomes/genbank/plant). For all three species, the genome fasta file, the genome annotation gff file and the complete CDS was obtained.

### 1) Determination of genomic nitrogen content

The genomic nitrogen content of each species was calculated by counting the number of occurrences of each nucleotide and multiplying by the corresponding number of nitrogen atoms using a python script and the genome fasta file. While a guanine-cytosine pair has eight nitrogen atoms, an adenine-thymine pair only has seven. The nitrogen content of the entire genome was determined first, then the nitrogen content of the CDS and non-CDS regions were also calculated using the CDS file.

### 2) Determination of transcriptomic nitrogen content and codon usage bias

The nitrogen content of all the pre-mRNA was determined by transcribing the regions annotated by ‘gene’ in the gff file using the Biopython (Bio.seq) library. Adenine has 5 nitrogen atoms, uracil has 2, guanine has 5 and cytosine has 3. The nitrogen content of the introns and exons were also determined separately. Finally, a codon usage table was obtained to examine preferential codon usage.

### 3) Determination of proteome nitrogen content

The nitrogen content of all the protein encoded by the CDS regions of each species was determined by counting the occurrences of each amino-acid and multiplying by the corresponding number of nitrogen atoms. The CDS sequences were converted to amino acid sequences using the Biopython library.

### 4) Determination of transfer-RNA nitrogen content and usage

tRNA genes were identified in each genome using the tRNAscan-SE software by Lowe, T.M. and Chan, P.P. (2016). tRNAscan-SE uses an advanced methodology for tRNA gene detection and functional prediction (determination of tRNA anticodon). Combining the results from the tRNAscan-SE and a python script, the nitrogen content of the tRNAs was determined. Using the data obtained from the codon usage tables obtained in part 2), it was possible to link codon biases with the corresponding tRNA nitrogen content. The aim was to examine whether tRNAs that had low nitrogen content had their corresponding codons more frequently represented than codons that were associated with higher nitrogen content tRNAs (among codons that are coding for the same amino acid).

## Results and Discussion

Investigating the nitrogen content of the genome of the three species reveals that *G. aurea* has a considerably lower number of nitrogen atoms in its genome than the two other plant species. A comparison with *T. pratense* shows the vast difference in nitrogen content can mostly be explained by the reduction of the number of nitrogen atoms in the non-coding DNA sequences of *G. aurea* (Fig. 1). Although *T. pratense* has 3 times more nitrogen in its CDS than *G. aurea*, the carnivorous plant has 10 times less nitrogen in its non-coding DNA sequences. At first glance, this observation supports the theory that genome reduction of *G. aurea* was motivated by nitrogen starvation. However, it might not come as a surprise to the reader that a vast reduction is genome size is accompanied by a vast reduction in genomic nitrogen content. This observation alone does not explain whether *G. aurea* has preferential usage of nitrogen-poor nucleotides (A-T base pairs).

**Figure 1:**
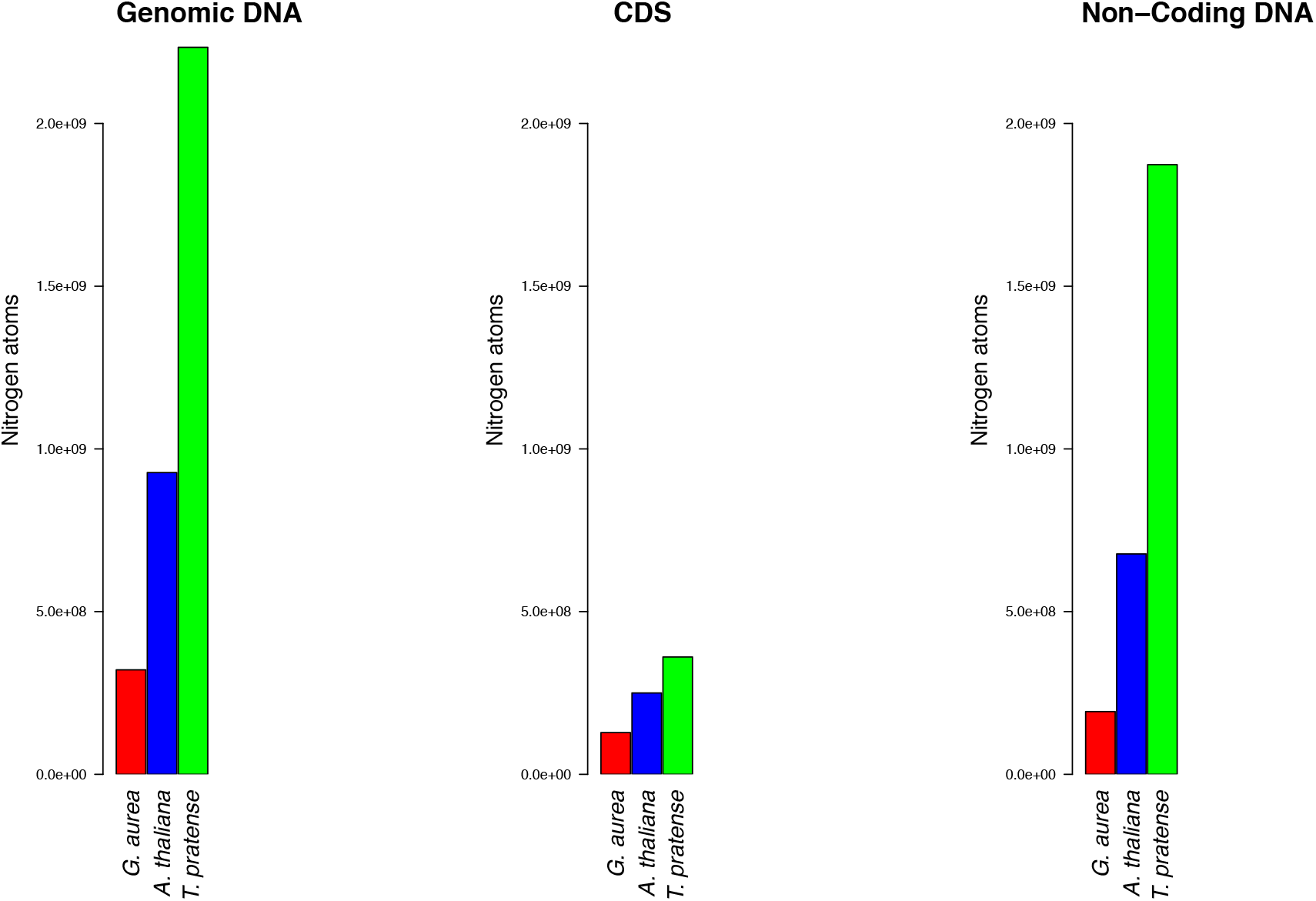
Number of nitrogen atoms in the entire genomic DNA, CDS and non-coding DNA of *G. aurea* (red), *A. thaliana* (blue) and *T. pratense* (green).

In this report, the term molecular unit refers to a DNA base-pair, an RNA nucleotide or a protein amino acid. Relative nitrogen content refers to the average number of nitrogen atoms per molecular unit. Upon examination of the relative nitrogen content of DNA, RNA and protein of the three-plant species, an unexpected pattern occurs (Fig. 2). The nitrogen starved carnivorous plant has higher nitrogen counts per molecular unit in genomic DNA, CDS, Non-Coding DNA, protein, mRNA, exons and introns. This data does not support the hypothesis that nitrogen starvation has caused preferential usage of molecular units that are lower in nitrogen. Inter species variations aside, CDS DNA was found to be higher in nitrogen than non-coding DNA and similarly exons were found to be higher in nitrogen than introns. Interestingly, in all 7 plots, *A. thaliana*, which has an intermediary genome length compared to *G. aurea* and *T. pratense*, was also found to have to have an intermediary nitrogen usage as well.

**Figure 2:**
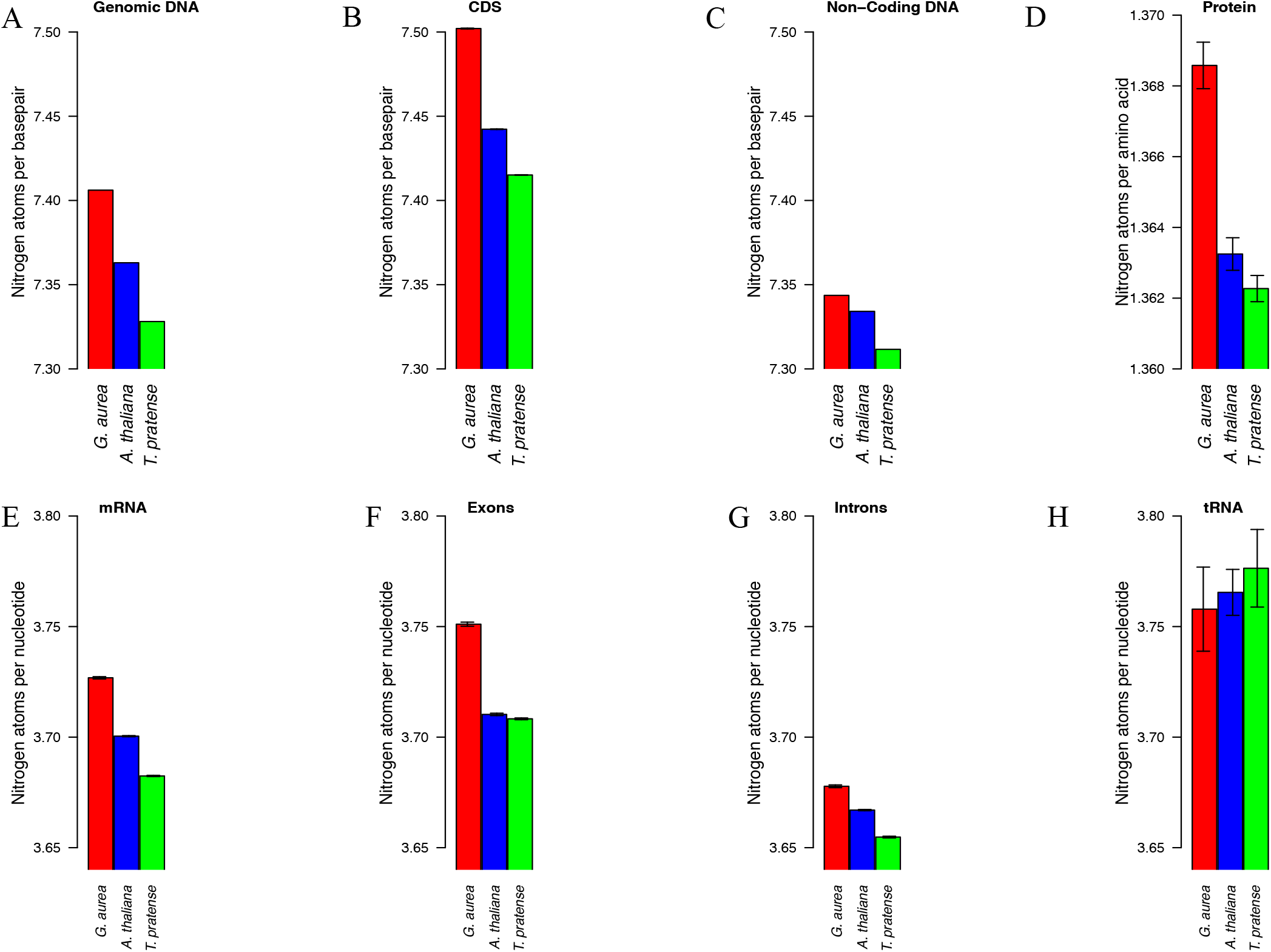
Average number of nitrogen atoms per molecular unit in genomic DNA, CDS, Non-Coding DNA, protein, mRNA, exons, introns and tRNA of *G. aurea* (red), *A. thaliana* (blue) and *T. pratense* (green). Error bars correspond to 95% confidence intervals. In this figure, mRNA refers to the pre-mRNA that hasn’t undergone removal of introns by splicing. Three different scales have been set for DNA sequences (A-C), protein sequences (D) and RNA sequences (E-H).

The first explanation of why *G. aurea* has a higher nitrogen usage in its DNA, RNA and proteins could be that there is not enough selective pressure on each molecular unit due to the small difference of nitrogen atoms gained for each molecular change. For example, a single substitution of a GC base-pair to an AT base-pair only lowers nitrogen usage by one nitrogen atom. In practice, it may be easier to remove whole sequences of non-coding or repeating sequences of DNA to optimize nitrogen usage. Some RNA transcripts may only be expressed for very short periods of time in the plant’s life cycle, reducing once again selective pressure on nitrogen optimization in these transcripts. However, this cannot explain why longer genomes are associated with lower nitrogen usage per molecular unit, at least for the three-species considered in this project. It is possible that the tiny genome of *G. aurea* combined with additional nitrogen captured from carnivory enables the species to have more leeway in using nitrogen rich amino acids and nucleotides. Finally, a last hypothesis is that G. aurea is actually using its transcriptome and proteome as a nitrogen bank. The high nitrogen content and the ubiquitous recycling of RNA and proteins in cells could make nitrogen storage in proteomes and transcriptomes possible.

Interestingly, the relative nitrogen content of tRNA may be lower in *G. aurea* than in the two-other species (Fig. 2, plot H). RNA sequencing data from Westermann, A., Gorski, S. and Vogel, J. (2012) shows that in eukaryotic cells there is about 3 times more tRNA than mRNA (as a measure of weight). The paper also states that cells contain almost 5 times more rRNA than tRNA, however, due to time constraints and the generally poorly annotated rRNA genes, rRNA nitrogen content was not determined. The fact that *G. aurea* had lower nitrogen content in tRNA sequences but not in other types of RNA or DNA sequences supports the hypothesis that there isn’t enough selective pressure on each molecular unit of DNA and mRNA to motivate nucleotide substitutions.

Studying the codon usage bias of *G. aurea* (Figure 3) revealed that the carnivorous plant uses the entire genetic code and this preliminary attempt to link codon usage bias with tRNA nitrogen concentration cannot conclusively draw a conclusion on whether codons complementary to tRNAs that are rich in nitrogen are less represented than codons complementary to tRNAs poor in nitrogen. For the majority of codons, multiple tRNA sequences were found to encode that codon, taking the mean of the nitrogen content of these sequences makes the assumption that all the tRNAs are equally expressed in *G. aurea*. This of course is extremely unlikely, thus sequencing the tRNA transcriptome of *G. aurea* is the only way to make this data more accurate.

**Figure 3:**
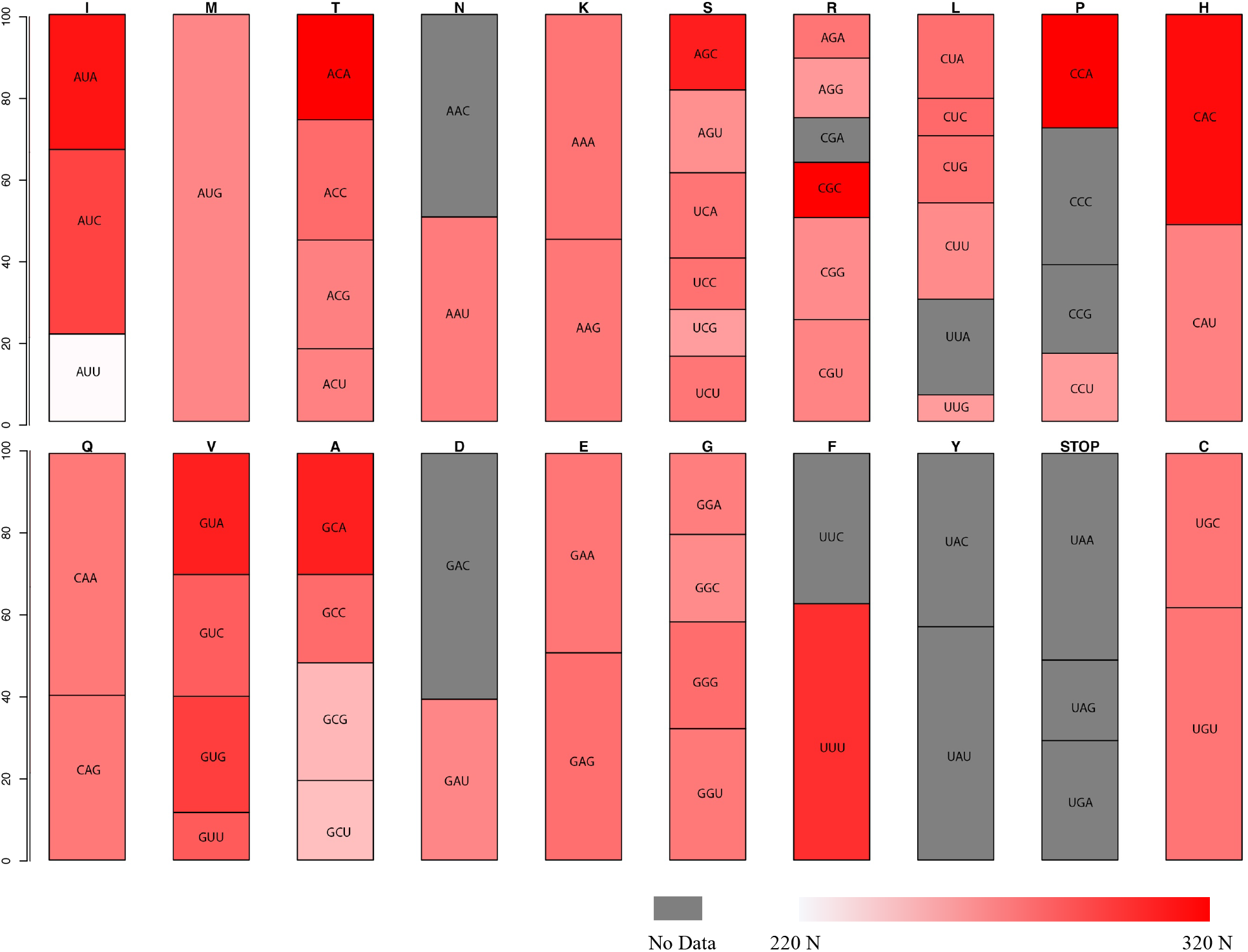
Bar graph representing the codon usage bias and tRNA nitrogen content in *G. aurea*. For each amino acid, the codon usage bias was determined and the relative proportion of each codon is represented. Codons that are complementary to tRNAs that are low in nitrogen are lighter in colour than codons complementary to tRNAs that are rich in nitrogen. When no tRNA sequences were found, the codon is represented in grey. When multiple tRNA sequences were found for a single codon, the average nitrogen of the tRNAs is represented. The colour scale bar ranges from 220 N atoms (pure white) to 320 N atoms (pure red). Note that Tryptophan (W) was removed for aesthetic reasons, no tRNA gene was found by tRNAscan-SE for tryptophan (grey).

## Conclusion

This comparative study of the carnivorous plant *Genlisea aurea*, the model organism *Arabidopsis thaliana* (Brassicaceae) and the nitrogen fixing *Trifolium pratense* (Fabaceae) attempted to investigate whether the genome, transcriptome and proteome of *G. aurea* showed evidence of adaptations to low nitrogen availability. It was found that although *G. aurea’s* genome, CDS and noncoding DNA were much lower in nitrogen than the genome of *T. pratense* and *A. thaliana* this was solely due to the length of the genome, CDS and non-coding sequences rather than the composition of these sequences. In fact, in the genomic DNA, CDS, non-coding DNA, mRNA, exons, introns and proteins of *G. aurea*, the relative nitrogen content was found to be greater than in the two-other species suggesting that nitrogen starvation might not put enough selective pressure on each molecular unit to motivate nucleotide substitutions. It was found that in tRNA sequences, which are about 5 times more abundant than mRNA in eukaryotes, *G. aurea* may have lower relative nitrogen. Finally, an attempt to link codon usage bias with the nitrogen content of complementary tRNAs proved inconclusive possibly due to the fact that multiple tRNAs can be complementary to a single codon. Future studies should determine the relative nitrogen content of ribosomal RNAs and perform transcriptome sequencing to determine the nitrogen content of the three species’ transcriptomes.

